# Determining the antiviral activity of two polyene macrolide antibiotics following treatment in Kidney Cells infected with SARS-CoV-2

**DOI:** 10.1101/2022.07.23.501242

**Authors:** Kishor M. Wasan, Chris Galliano

## Abstract

**Background:** There has been much speculation that polyene macrolide antibiotics, such as amphotericin B (AmB) and Nystatin (NYS) may have antiviral activity against several viruses including SARS-CoV-2.

**Objective:** The objective of this short communication was to determine the antiviral activity of two polyene macrolides, AmB and NYS, following treatment in kidney cells infected with SARS-CoV-2.

**Methods:** A serial dilution of AmB, NYS, and irbesartan (a drug known to bind to the ACE-2 receptor as a positive control) were then added (n=4 at each concentration) to the infected Vero’76 kidney cells in 100 µL media. Cells were also examined for contamination at 24 hours, and for cytopathic effect (CPE) and cytotoxicity (if noticeable) under a microscope at 48 hours. In a second study, AmB and Remdesivir were incubated in kidney cells infected with the virus and inhibition of the virus was determined by an immunoassay.

**Results and Conclusions:** Amphotericin B (AmB) showed a significant reduction in the TCID50 titer, with the 50% effective concentration (EC50) of 1.24 µM, which was 2.5 times lower than the cytotoxicity concentration. NYS and Irbesartan both exhibited substantially less active and would not be considered a suitable choice for further investigations. In addition, when measuring viral inhibition by immunoassay, AmB was significantly more potent than remdesivir (EC50 31.8 nM vs. 1.15 µM). Taken together, these preliminary findings suggest that AmB may have significant activity against SARS-CoV-2. However, further cell and animal studies are warranted.

## 1. Introduction

The emergence of COVID-19 has affected every population in the world causing millions of infections. To date, there are very few therapeutics to treat patients with COVID-19 that exhibit mild or moderate symptoms but not require hospitalization. The development of an effective, safe, and proven vaccine is clearly the key step in controlling the pandemic, however, there is still a need for effective therapeutics to treat those infected with the virus. Amphotericin B (AmB) is a polyene macrolide antibiotic used for the treatment of systemic fungal and parasitic infections [1-10]. There has been much speculation that AmB may have antiviral activity against several viruses including the ones responsible for COVID-19 [11-15]. Recent investigations have suggested that AmB may interact with the S-protein of the SARS-CoV-2 virus, thus blocking its interaction with the ACE-2 receptor found on epithelial and endothelial cells [11-15]. Thus, the hypothesis of this study was that AmB will function as an antiviral not only by binding to and preventing SARS-CoV-2 virus replication, but also by blocking viral uptake into mammalian cells. Our laboratory tested this hypothesis by determining the antiviral activity of AmB against SARS-CoV-2 within kidney cells.

## 2. Materials and Methods

Vero’76 (ATCC CRL-1587) kidney cells were seeded and grown overnight at 37□ in a 5% CO2 environment to approximately 90% confluence in 96-well plates. The culture medium used was DMEM supplemented with 10% FBS/PEN-STREP. The virus SARS-CoV-2/Canada/ON/VIDO-01/2020/Vero’76/p.2 (Seq. available at GISAID – EPI_ISL_413015) was diluted in DMEM supplemented with 2% FBS/PEN-STREP to obtain a multiplicity of infection (MOI) of 0.1 (approximately 2000 TCID_50_/well). The culture medium was removed from the cells and 50 µl of virus inoculum was added to each well. The plates were incubated for 1 hour at 37□ in a 5% CO_2_ environment, then the medium was removed. A serial dilution of AmB, NYS, and Irbesartan (a drug known to bind to the ACE-2 receptor as a positive control) were then added (n=4 at each concentration; data presented as mean +/- SD; SD are very small of the size of the graph) to the infected cells in 100 µL media. Medium alone was used as a control for virus replication, and untreated, uninfected control wells were also established. A separate plate was used to assess the cell toxicity of the test compounds by incubating uninfected cells with the serially diluted test compound solution. After 48 hours of incubation, cytotoxicity was measured by MTS assay (CellTiter 96 Aqueous One (Promega. Spectroscopic absorbance at 490 nM following incubation is directly related to cytotoxicity. The corrected absorbance at 490 nM versus the concentration of AmB, NYS, and irbesartan (n=4 at each concentration) was plotted to determine CC50, the concentration at which 50% of the cells were dead. Cells were also examined for contamination at 24 hours, and for cytopathic effect (CPE) and cytotoxicity (if noticeable) under a microscope at 48 hours. At 48 hours, 100 µl supernatant from each well was harvested. Viral titration by the TCID_50_ assay of the supernatant was carried out by a serial 10-fold dilution. These dilutions were subsequently used to infect another set of plated cells, in quadruplicates as described for the initial infection. Cells were observed for CPE at 1, 3, and 5 days after infection.

In a second set of studies, efficacies were tested in parallel in African green monkey kidney (Vero E6) cells. Each test compound was tested individually and each of the concentrations was evaluated in triplicate for efficacy. Vero E6 cells were cultured in 96 plates prior to the day of the assay. Cells were greater than 90% confluency at the start of the study. Each of the test article concentrations was evaluated in triplicate. Test article concentration were tested in two different conditions: 1) Pre-treatment for 24 ± 4 hours prior to virus inoculation followed by treatment immediately after removal of the virus inoculum or 2) treatment only with the test article added immediately following removal of the virus inoculum. These two different conditions were tested to ascertain the effect of the drugs on the entry, fusion, replication, and transmission of the virus.

Remdesivir was added immediately following removal of the virus inoculum. For pre-treatment and treatment, wells were overlaid with 0.2 mL DMEM2 (Dulbecco’s Modified Eagle Media (DMEM) with 2% Fetal Bovine Serum (FBS) to test articles at concentrations as delineated in Section 9.8). Following the 24 ± 4 hour pretreatment, cells were inoculated at a MOI of 0.001 TCID50/cell with SARS-CoV-2 and incubated for 60-90 minutes. Immediately following the 60–90-minute incubation, the virus inoculum was removed, cells washed, and appropriate wells overlaid with 0.2 mL DMEM2 (DMEM with 2% FBS with test or control articles) and incubated in a humidified chamber at 37°C ± 2°C in 5 ± 2% CO_2_. At 48 ± 6 hours post inoculation, cells were fixed and evaluated for the presence of virus by immunostaining assay justification: The immunostaining assay utilized a modified incubation time to 48 hours. A 24 ± 4-hour pre-treatment of the cells was included for selected test articles. After 48 ± 6 hours, cells are fixed with paraformaldehyde and stained by anti-SARS-2 nucleoprotein monoclonal antibody (Sino Biological) followed by peroxidase-conjugated goat anti-mouse IgG (SeraCare). Wells are developed using TMB Substrate Solution and the reaction stopped by acidification. The ELISA plate were read at 450 nm on a spectrophotometer by ELISA plate reader.

For each well, the inhibition of virus is calculated as the percentage of reduction of the absorbance value in respect to the virus control by the following formula: percent inhibition = 100 - [(A450 of article dilution - A450 of cell control)/(A450 of virus control - A450 of cell control)] x 100. The EC50 is defined as the reciprocal dilution that caused a 50% reduction of the absorbance value of the virus control (50% A450 reduction).

## 3. Results

Amphotericin B (AmB) showed a significant reduction in the TCID50 titer, with the 50% effective concentration (EC50) of 1.24 µM, which was 2.5 times lower than the cytotoxicity concentration (**Table 1**). Nystatin (NYS), a compound similar in chemical structure to Amphotericin B and irbesartan both exhibited substantially less active and would not be considered a suitable choice for further investigations. Figures 1 through 3 demonstrate the difference in concentration response of NYS and irbesartan compared to AmB, respectively, where AmB exhibits a 32-fold lower concentration required to achieve EC50 at a 15-fold lower CC50 compared to nystatin. The selectivity index (SI) was calculated as the ratio of CC50 to EC50. The SI for both compounds was > 1, which indicates that the compounds are more effective than they are toxic to the cells (**Table 1**).

**Table 1.** Vero-76 Kidney Cells Infected Cells (Viral Strain: (SARS-CoV-2/Canada/ON/VIDO-01/2020/Vero’76/p.2)* treated with Amphotericin B, Nystatin and Irbesartan.

| Drug | EC<sub>50</sub> | SI |
| --- | --- | --- |
| Amphotericin B | 1.24 μM ** | 3.1 |
| Nystatin | 36.6 μM | 1.41 |
| Irbesartan | 62.5 μM | 5.55 |

\*D614G variant that has been circulating across the continent since early 2020 and there have been no appreciable differences in transmission or virulence across cities in North America that would suggest any mutational changes in the virus resulting in a new variant. \*\*Activity based on concentrations used to treat systemic fungal infections (1000-2000 nM).

**Figure 1.**
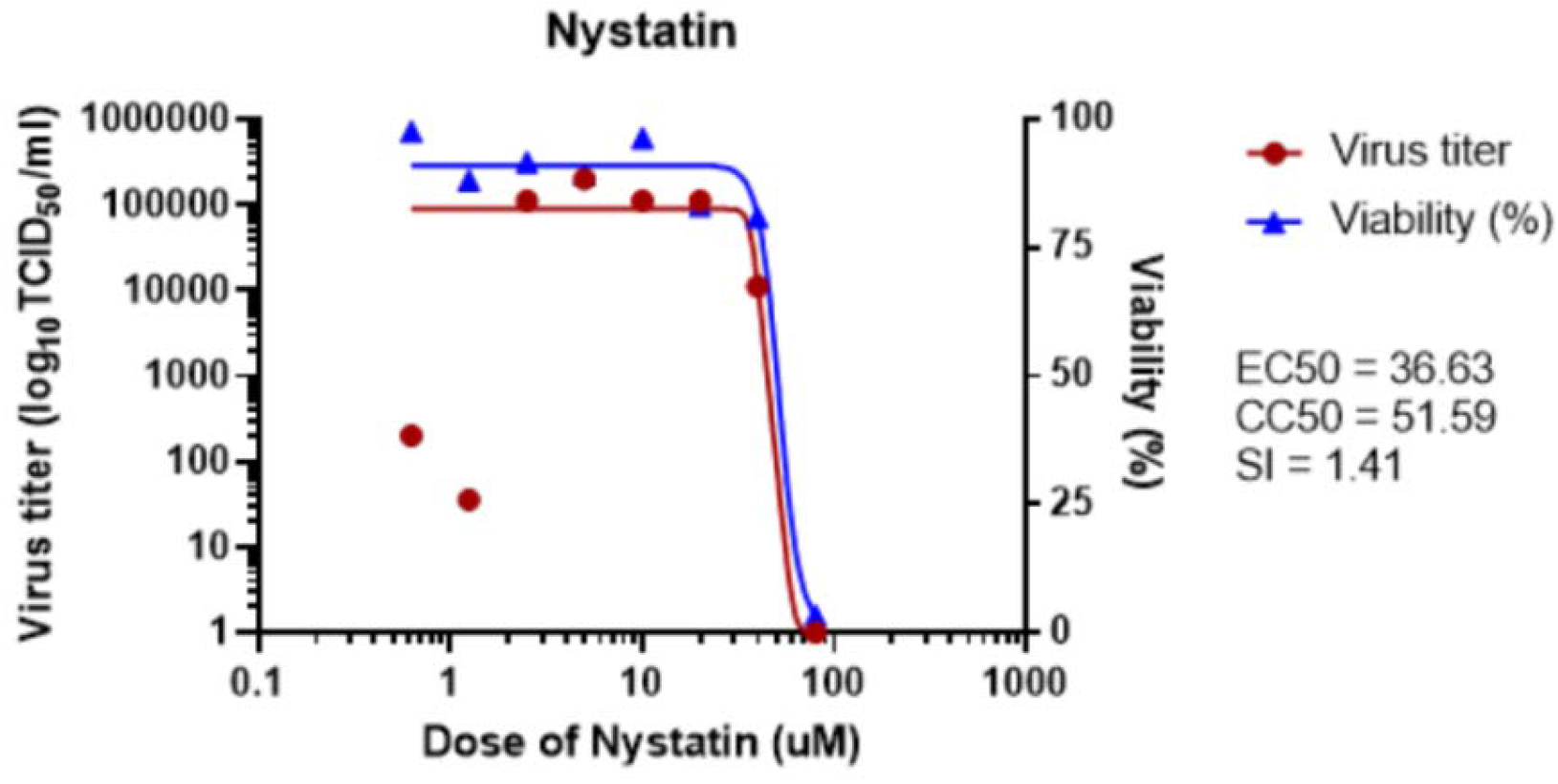
Virus titer and cell viability following increasing concentrations of nystatin have been incubated with Vero kidney cells for 48 hours that have been infected with the SARS-CoV-2 virus. Data is presented as mean +/- standard deviation (n=4 for each concentration tested).

**Figure 2.**
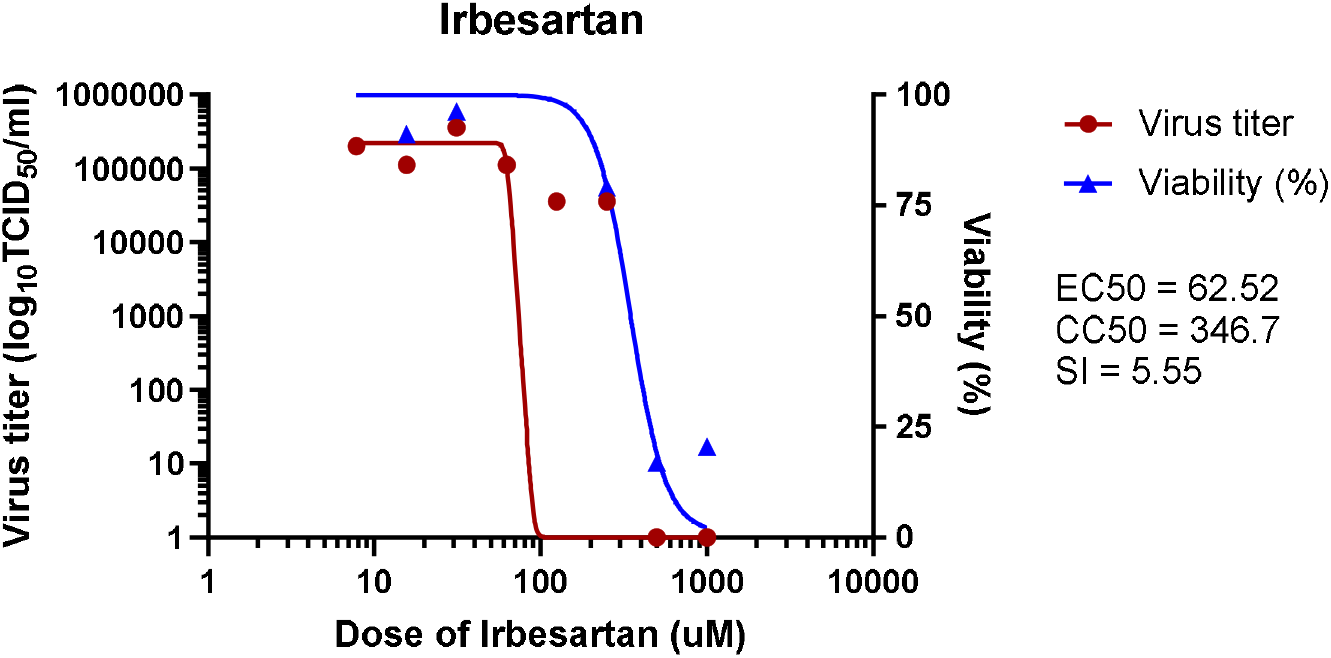
Virus titer and cell viability following increasing concentrations of irbesartan have been incubated with Vero kidney cells for 48 hours that have been infected with the SARS-CoV-2 virus. Data is presented as mean +/- standard deviation (n=4 for each concentration tested).

**Figure 3.**
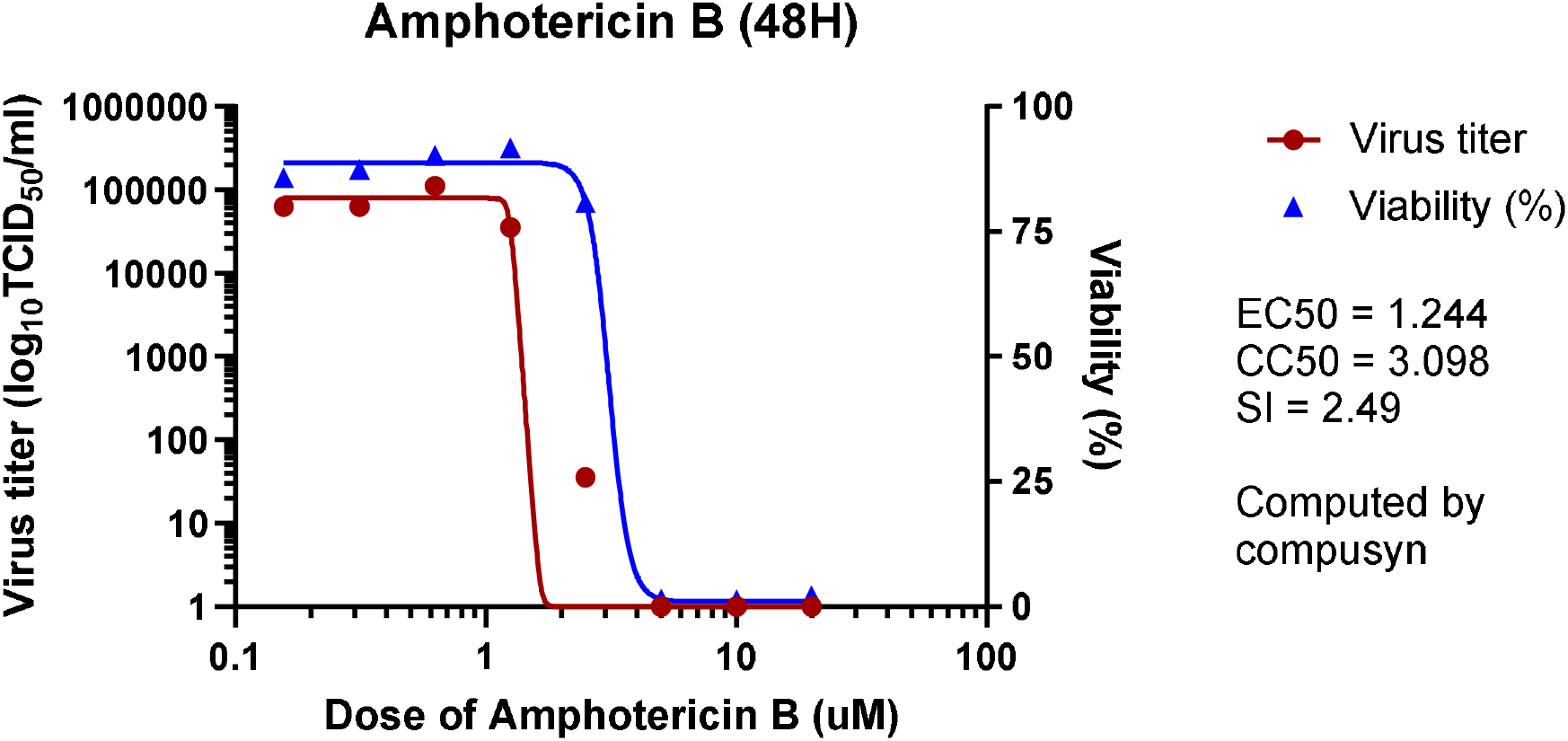
Virus titer and cell viability following increasing concentrations of Amphotericin B have been incubated with Vero kidney cells for 48 hours that have been infected with the SARS-CoV-2 virus. Data is presented as mean +/- standard deviation (n=4 for each concentration tested).

The above results of **Table 2** confirm the results obtained in the first testing series and confirm the versatility of AmB as a replication blocker of COVID-19 as the tests were carried out on different strains of COVID-19. The above results also indicate that in this test, AmB was clearly superior to remdesevir in terms of EC50.

**Table 2.** Efficacy in African green monkey kidney (Vero E6) cells infected with Virus (Viral Strain :2019 Novel Coronavirus, Isolate USA-WA1/2020 (SARS-CoV-2)Code.

| Drug | EC<sub>50</sub> | EC<sub>100</sub> | comment |
| --- | --- | --- | --- |
| Amphotericin B | 31.8 nM** | Achieved |  |
| Remdesevir | 1.15um | Achieved | Positive control |

\*\*Activity based on concentrations used to treat systemic fungal infections (1000-2000 nM).

## 4. Discussion and Conclusions

Taken together, these preliminary findings suggest that AmB may have significant activity against SARS CoV2 and may be efficacious in patients suffering from COVID-19 where better treatment is still needed. Further cell and animal studies are required to confirm these important preliminary findings and could lead to the development and implementation of AmB in the treatment of COVID 19.

## Author Contributions

KM and CG were all involved in the conceptualization, methodology and writing—original draft preparation and writing—review and editing of this manuscript. All authors have read and agreed to the published version of the manuscript.

## Funding

Skymount Medical provided funding for this research. We also received in-kind support from VIDO-InterVac at the University of Saskatchewan that completed some this preliminary work in their BCL-3 facility.

## Acknowledgments

We want to thank VIDO-InterVac at the University of Saskatchewan for their support.

## Conflicts of Interest

Declare conflicts of interest or state “The authors declare no conflict of interest.”

## Notes

### Competing Interest Statement

The authors have declared no competing interest.

